# The genome of the poecilogonous annelid *Streblospio benedicti*

**DOI:** 10.1101/2021.04.15.440069

**Authors:** Christina Zakas, Nathan D. Harry, Elizabeth H. Scholl, Matthew V. Rockman

## Abstract

*Streblospio benedicti* is a common marine annelid that has become an important model for developmental evolution. It is the only known example of poecilogony, where two distinct developmental modes occur within a single species, that is due to a heritable difference in egg size. The dimorphic developmental programs and life-histories exhibited in this species depend on differences within the genome, making it an optimal model for understanding the genomic basis of developmental divergence. Studies using *S. benedicti* have begun to uncover the genetic and genomic principles that underlie developmental uncoupling, but until now they have been limited by the lack of availability of genomic tools. Here we present an annotated chromosomal-level genome assembly of *S. benedicti* generated from a combination of Illumina reads, Nanopore long reads, Chicago and Hi-C chromatin interaction sequencing, and a genetic map from experimental crosses. At 701.4 Mb, the *S. benedicti* genome is the largest annelid genome to date that has been assembled to chromosomal scaffolds, yet it does not show evidence of extensive gene family expansion, but rather longer intergenic regions. The complete genome of *S. benedicti* is valuable for functional genomic analyses of development and evolution, as well as phylogenetic comparison within the Annelida and the Lophotrochozoa. Despite having two developmental modes, there is no evidence of genome duplication or substantial gene number expansions. Instead, lineage specific repeats account for much of the expansion of this genome compared to other annelids.

## Introduction

The annelids, which are grouped within the Lophotrochozoa, are a large but generally understudied animal lineage in terms of development and genomics. Genomic sequencing of Lophotrochozoan animals has yielded important discoveries in genome evolution, adaptation, and novelty (Simakov et al. 2013; Albertin et al. 2015; Schiemann et al. 2017; Wang et al. 2017; Luo et al. 2018).Conserved genes and pathways across the Bilateria have been investigated in other animal branches but may have novel functions and divergent outcomes in these groups that have yet to be explored (Tessmar-Raible and Arendt 2003; Paps et al. 2015). The Lophotrochozoa are a superb group for studying development, adaptation, and evolution on the genomic level due to the conserved patterns of ontogeny and a unique range of novel adaptations (Henry and Martindale 1999; Seaver 2014). However, they are severely lacking genomic resources: currently nine full genome datasets are listed in NCBI for Annelids. Despite the biodiversity contained in this group, little is known about genomic evolution and how that informs important process like convergence, gene regulatory network modification, and developmental systems drift. This gap in genomic resources impairs our understanding of basic developmental and evolutionary biology in a major animal lineage.

The marine annelid *Streblospio benedicti* (Figure 1) is a particularly important model for understanding the genomic basis for developmental evolution. *S. benedicti* is one of the rare cases of confirmed *poecilogony* - two distinct modes of development occurring within the species. *S. benedicti* exhibits both indirect development, with a distinct larval phase, and direct development, with offspring that resemble small adult forms (Levin 1984; Levin and Bridges 1994). In *S. benedicti* adults are gonochoristic (separate males and females) and females produce a fixed offspring type throughout their lives (Levin et al. 1987). There are two types of females in *S. benedicti*, which are essentially indistinguishable other than traits related to producing offspring (Gibson et al. 2010). The two types of females differ in the egg sizes they produce (~100 versus ~200 μm diameter eggs) and per-clutch fecundity (~200-400 versus ~20-50 offspring per clutch; McCain 2008). The resulting offspring are drastically different, with contrasting development modes (*planktotrophy*: obligately feeding larvae, versus *lecithotrophy*: non-obligately-feeding larvae). These larval types differ in their planktonic development time (2-4 weeks swimming in the plankton versus 0-2 days before settlement), larval ecologies (pelagic versus benthic larvae) and overall life-history strategies. Importantly, these trade-offs only occur during the embryonic and larval phases; by the time the worms become adults they are morphologically indistinct aside from some female reproductive anatomy (number of brood pouches and segments on which the brood pouches occur) and they occupy the same types of estuarine environments (Gibson et al. 2010). Interestingly, there is gene flow between adults of different types, but they usually do not directly co-occur (Zakas and Wares 2012). Furthermore, *S. benedicti* is the only known case of poecilogony where the developmental types are heritable with a strong additive genetic basis, as opposed to plasticity or polyphenism (Levin et al. 1991; Zakas and Rockman 2014; Zakas et al. 2018). Because these differences in development and life-history are contained within a single species, the ability to investigate the genomic basis of developmental variation is unparalleled. The genome of *S. benedicti* provides an opportunity to explore how a major transition in animal development happens on the genomic level.

**Figure 1.**
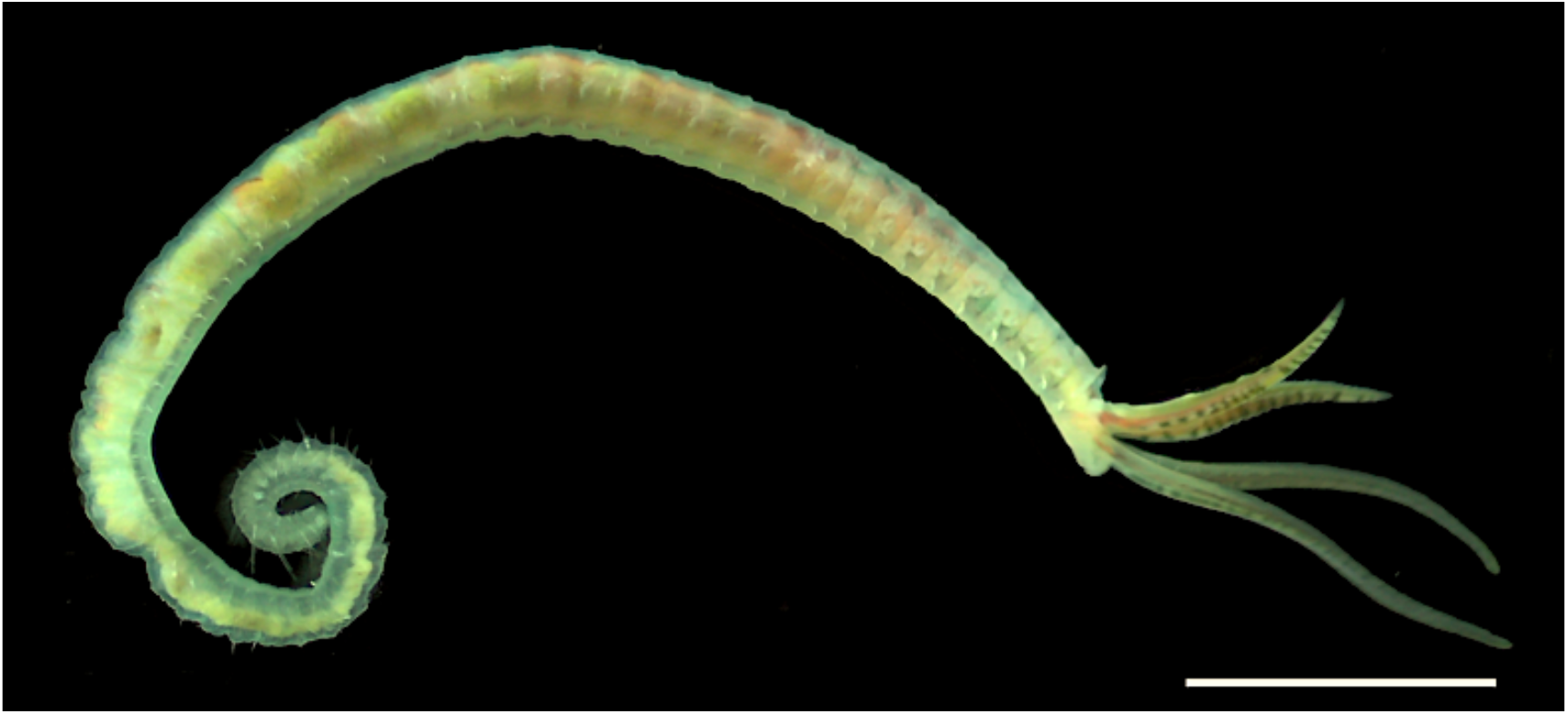
Adult *S. benedicti*. Scale bar is 1mm.

High-quality genome assemblies are now enabling technical advances in using new models with naturally occurring traits of interest such as poecilogony. Here we present the genome of *S. benedicti*, which we constructed from a combination of Illumina short reads and Nanopore long reads, scaffolded with Hi-C and Chicago proximity-ligation data. We use the previously described genetic linkage map of *S. benedicti* to correct scaffolding arrangements and locate quantitative trait loci (QTL) markers in the genome (Zakas et al. 2018). This is the one of the few high-quality annelid genomes and opens opportunities for transformative research in this and related systems.

## Results

### Genome Sequencing

We assembled the chromosomal-level genome of *S. benedicti* using individuals of the planktotrophic morph from Bayonne, New Jersey. The genome is assembled from five classes of data:

1. PCR-free Illumina shotgun sequence data generated from a pool of 13 females from a single F_9_ inbred line.
2. Nanopore long-read data generated from multiple pools including 71 wild-caught females
3. Chicago proximity-ligation data generated from a pool of three males from the inbred line.
4. Hi-C data generated from a pool of two females from the inbred line.
5. Genetic map data from an experimental cross

We recovered between 6-8.5 million reads per Nanopore flow cell, and 3-5 million reads were over 6.5 Kb long. In total we generated 30 million reads (with 15.6 million >6.5 Kb). Illumina shotgun data alone yielded a draft assembly with scaffold N50 of 53 Kb. The addition of the Nanopore, Chicago, and Hi-C data improved the assembly by an order of magnitude: increasing the scaffold N50 to 53.55 Mb (Table 2, Figure S1).

**Table 1.** Scaffold length and names for 11 chromosomes.

| Linkage Group | Scaffold Name | Scaffold length (Mb) |
| --- | --- | --- |
| Chr1 | Sc8CjP1_6019 | 38.1 |
| Chr2 | Sc8CjP1_6202 | 62.6 |
| Chr3 | Sc8CjP1_6205 | 47.2 |
| Chr4 | Sc8CjP1_6198 | 53.9 |
| Chr5 | Sc8CjP1_5318 | 56.1 |
| Chr6 | Sc8CjP1_6207 | 64.9 |
| Chr7 | Sc8CjP1_32 | 46 |
| Chr8 | Sc8CjP1_5307 | 64.8 |
| Chr9 | Sc8CjP1_2914 | 61.6 |
| Chr10 | Sc8CjP1_78 | 43.7 |
| Chr11 (X) | Sc8CjP1_6176 | 61.4 |
| Total Genome in Chromosomes |  | 600.3 (85.5%) |
| Total Genome in unplaced scaffolds |  | 101.2 (14.5%) |

**Table 2.** Summary statistics of genome assembly.

|  | **Dovetail** | **Chromonomer** |
| --- | --- | --- |
| Assembly Size | 702.4 Mb | 701.4 Mb<sup>b</sup> |
| Heterozygosity (SNP%) | 0.29 (31-mers) | - |
| Number of scaffolds | 6207 | 6112 |
| L50/N50 | 6/ 53.3 Mb | 6/ 56.1 Mb |
| Total gene models predicted | 20,767 | 20,221 |
| BUSCO score<sup>a</sup> | 84.5(S) 2.3(D) 5.3(F) 7.9(M) | 88.4(S) 3.4(D) 2.4(F) 5.8(M) |
| BUSCO total | 92.1 | 94.2 |
| GC% | 37.8 | 37.8 |
| %N | - | 0.3 |

<sup>a</sup>(BUSCO: S=single, D= duplicated, F=fragmented, M=missing)
<sup>b</sup>size reduction from contamination removal

### Genome Assembly

To integrate the genetic map with the reference genome, we corrected the potential mis-assembly of scaffolds using Chromonomer (Catchen et al. 2020). Chromonomer breaks super-scaffolds in regions of low confidence (stretches of ‘N’ ambiguity) and rearranges them based on high-confidence markers in the genetic map. The *S. benedicti* genetic map was previously constructed from G2 families with 702 markers in 11 linkage groups (Zakas et al. 2018). Chromonomer used 570 (81% of the total) informative markers to reconcile the reference to the genetic map. The resulting assembly increased the genome’s N50 by 2.8 Mb, reduced the number of scaffolds in the assembly, and increased its BUSCO (Benchmarking Universal Single-Copy Orthologs) score by combining many smaller scaffolds with chromosomal scaffolds (Table 2). There are 11 chromosome-level scaffolds, from 38-65 Mb in length. They correspond to the karyotype, which shows ten autosomes and one sex chromosome, and the 11 linkage groups constructed previously (Zakas et al. 2018).

### Gene Annotations

An Iso-Seq transcriptome of pooled planktotrophic individuals of all developmental stages was used for annotations. Based on GMAP (version 2020-06-30; Wu and Watanabe 2005) 24,117 of 24,317 high-quality Iso-seq transcripts mapped uniquely to the genome.

Using the Iso-Seq data as well as proteins from *Capitella teleta* as evidence, Maker v2.31.10 called 41,088 transcripts across the entire genome, including at least one gene on 1,899 of the 6,101 unplaced scaffolds. These transcripts represent 20,221 genes, indicating an average of 2 transcripts per locus throughout the genome. Additionally, 5,995 tRNA were identified (although 3,821 are noncoding/pseudo tRNAs).

### Repeat Modeling

The genome of *S. benedicti* is 40.36% repetitive, which is greater than other annelids reported (Table 3, Table S4), and it contains substantially more interspersed repeats. Interestingly, most repeats are unclassified (comprising 30.19% of the genome) and are likely lineage-specific elements. This is not particularly unusual as emerging models tend to have novelity in repetitive elements and the families of repeats in the Lophotrochozoa remain understudied.

**Table 3.** Gene summary statistics.

|  |  |
| --- | --- |
| Average gene length | 3.54 Kb |
| Median exon length | 157 bases |
| Median intron length | 410 bases |
| Transcribed regions | 17% |
| Mean number of exons per transcript | 3.1 |
| Protein-coding genes | 20,221 |
| mRNAs | 41,088 |
| Exons | 125,713 |
| CDSs | 124,795 |
| 5' UTRs | 8,260 |
| tRNAs | 5,995 |
| 3' prime UTRs | 6,833 |

### Comparison with other Annelid Genomes

We compared the *S. benedicti* genome to three other annelid reference genomes: *Capitella teleta*, which is the most closely related species with a reference genome, *Helobdella robusta* (leech) *and Dimorphilus gyrociliatus*, which is the most compact and complete sequenced annelid genome (Simakov et al. 2013; Martín-Durán et al. 2020). *S. benedicti* has the largest and least gene-dense genome. There are fewer genes per megabase and more interspersed repeats in *S. benedicti* (Table 4).

**Table 4.** Annelid genome comparisons.

|  | *Capitella teleta* | *Helobdella robusta* | *Dimorphilus gyrocilatus* | *Streblospio benedicti* |
| --- | --- | --- | --- | --- |
| Genome Size | 334 Mb | 235 Mb | 77.9 Mb | 701.4 Mb |
| Scaffold Number | 21,042 | 1,993 | 349 | 6,112 |
| Scaffold N50 | 188 Kb | 3.06 Mb | 2.24 Mb | 56.1 Mb |
| Longest Scaffold | 1.62 Mb | 13.6 Mb | 5.53 Mb | 64.8 Mb |
| Unknown Sequences | 17.0% | 8.5% | 1.7% | 0.3% |
| BUSCO | 96.6% | 89.6% | 95.8% | 94.2% |
| Genes per Mb | 100.0 | 97.5 | 208.9 | 28.8 |
| Median Intron Length | 57 | 255 | 66 | 410 |
| Unique Proteins | 29% | 21% | 17% | 27% |
| Protein Coding Genes | 32,175 | 23,432 | 14,203 | 20,221 |
| Unique Clusters | 1794 | 727 | 759 | 1948 |
| Enriched GOterms | 8 | 0 | 0 | 28 |
| Total Masked | 31.06% (103 Mb) | 33.39% (78 Mb) | 11.18% (8.71 Mb) | 40.36% (283 Mb) |
| Interspersed Repeats | 22.87 (76.39 Mb) | 18.46% (43.38 Mb) | 7.95% (6.19 Mb) | 39.16% (275 Mb) |
| Simple Repeats | 3.68% (12.3 Mb) | 12.21% (28.69 Mb) | 2.69% (2.09 Mb) | 1.10% (77 Mb) |
| Low Complexity Repeats | 4.24% (12.15 Mb) | 2.58% (6.07 Mb) | 0.55% (0.43 Mb) | 0.13% (0.87 Mb) |

We used OrthoVenn2 (Xu et al. 2019) to find gene cluster distributions across the four genomes, based on our dataset of 41,088 predicted transcripts. Gene clusters contain sets of orthologs or paralogs, and overlapping clusters contain proteins from different species. There are 15,259 clusters in this comparison and 8,511 of these contain *S. benedicti* proteins (Figure 2). There are 1,948 (23% of all *S. benedicti* clusters) that are paralog clusters unique to *S. benedicti*, which is more than the other annelids. But unique proteins make up only 27% of the *S. benedicti* total proteins, which is comparable to the other species (Table 4). There are 11,281 transcripts found only in *S. benedicti* including those in the 1,948 clusters as well as 6,508 genes that are single-copy and single-isoform and are therefore not part of any clusters (Figure S2). *S. benedicti* has more predicted transcripts than the other annelids, but a similar number (20,211) of protein coding genes and a similar number of total gene clusters (Figure 2, Table S1). The larger number of transcripts found in the *S. benedicti* genome reflects an average two transcripts per gene although the distribution varies (Figure 3). More robust comparisons about genomes and gene expansion can be addressed with the addition of new and updated Annelid genomes available in the near future.

**Figure 2.**
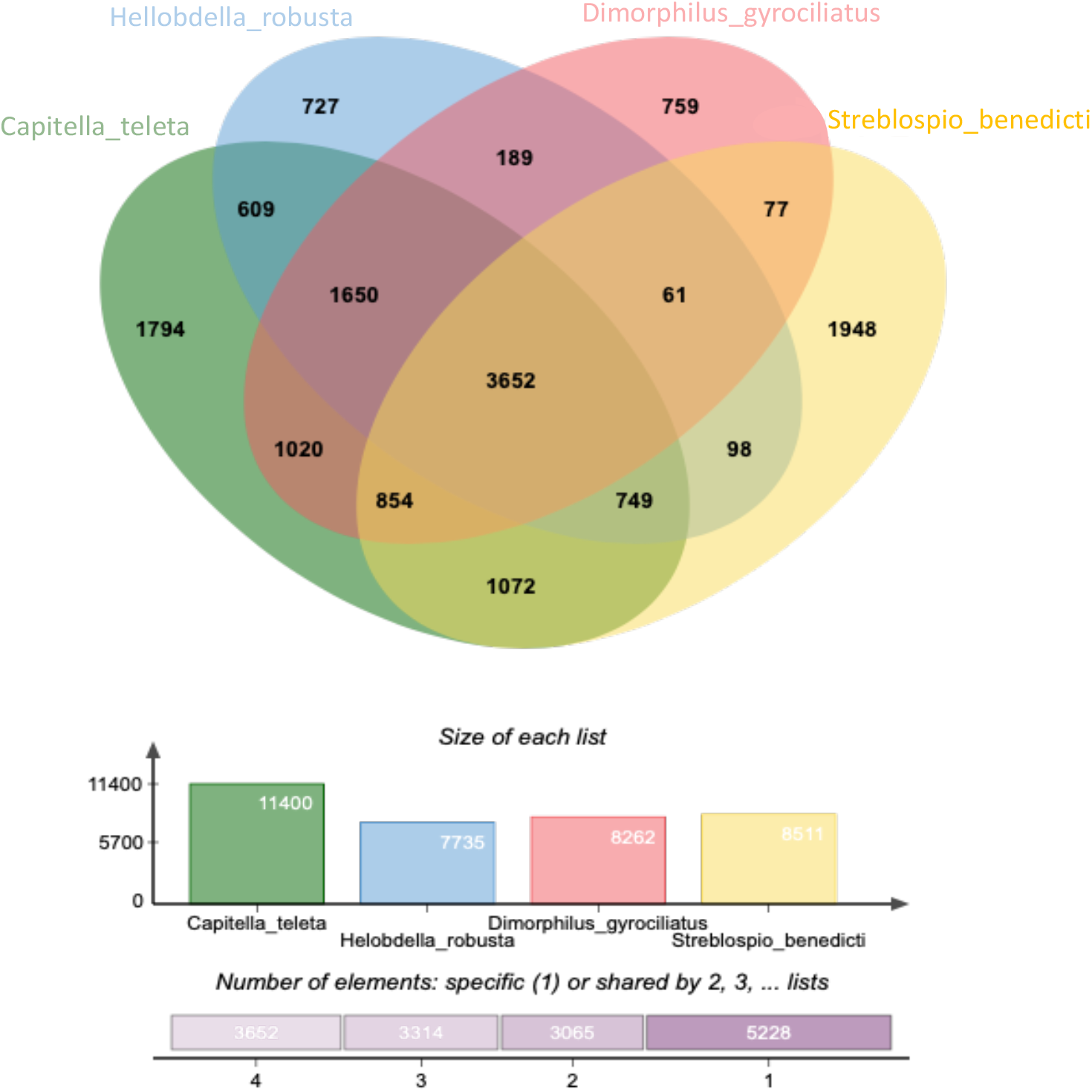
Comparison of four annelid genome ortholog clusters.

**Figure 3.**
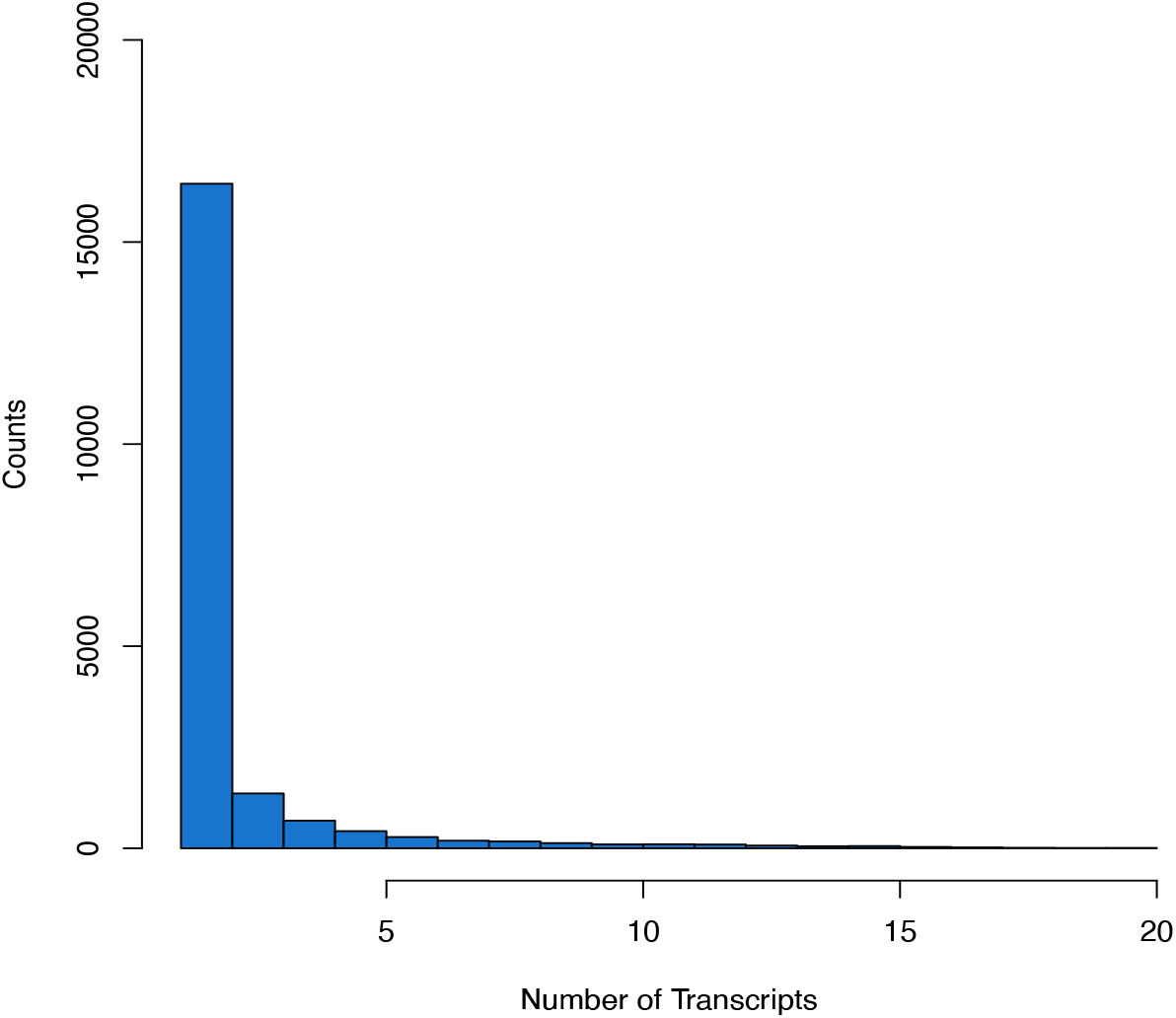
Histogram of transcripts per gene.

OrthoVen2 assigned GO terms to each of the gene clusters. Of the clusters unique to *H. robusta* (727) and *D. gyrociliatus* (759) there are no enriched GO terms relative to the other groups (Table 4). In *C. teleta*’s unique gene clusters (1794) there are 8 GO terms that are enriched, while In *S. benedicti* there are 28 enriched GO terms listed in Table S2, Figure 4. The proportion of unique clusters, singletons, and enriched GO categories suggest that there are more novel genes in the *S. benedicti* genome compared to the other annelids, although there is a range of novel genes reported across taxa within the Lophotrochozoa (Sun et al. 2020) and novel gene clusters are not necessarily correlated with functional novelty.

**Figure 4.**
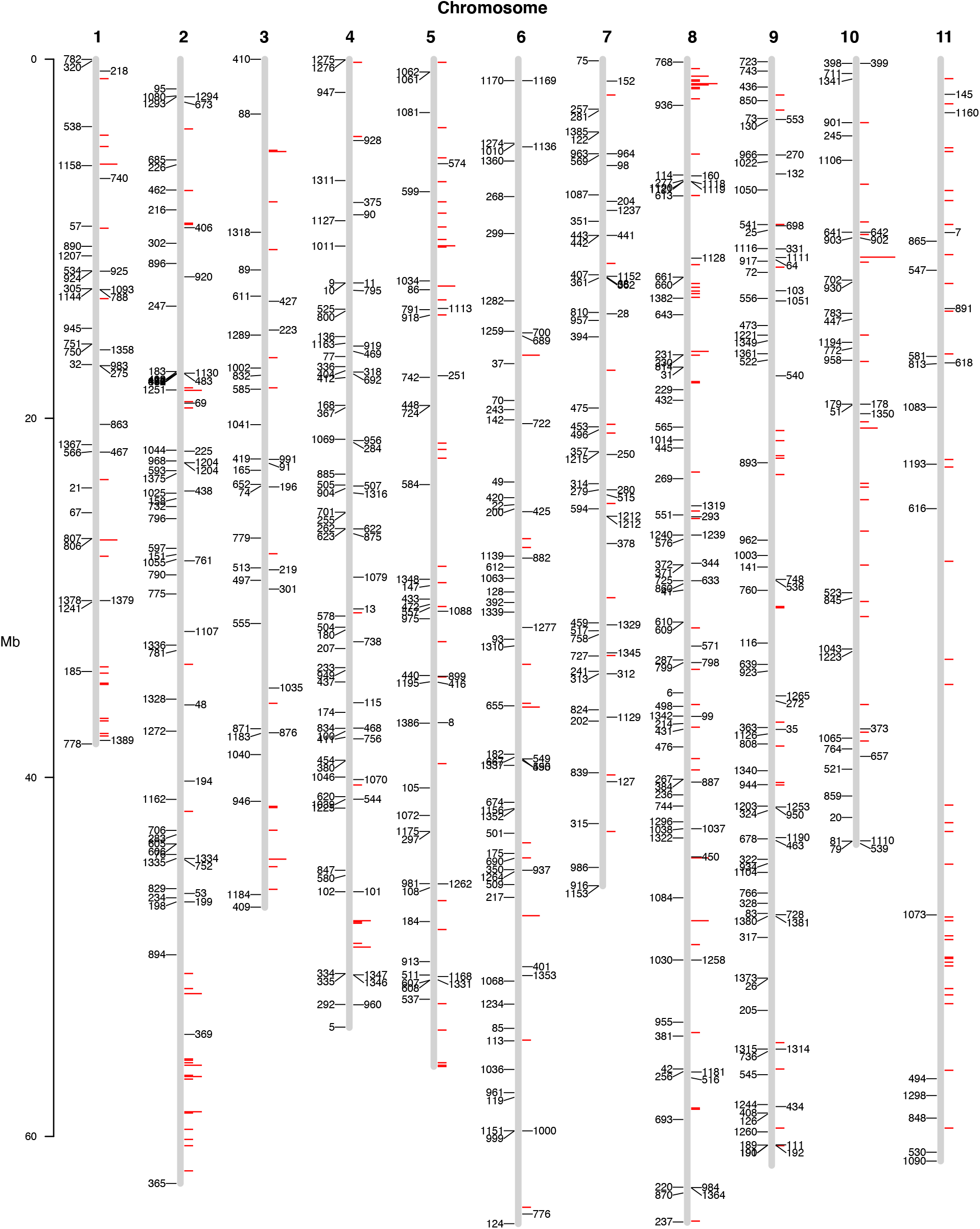
Plot of genome assembly for 11 chromosomes. Red histogram shows the placement of gene transcripts identified in the *S. benedicti* GO enrichment categories. 595 of 702 linkage markers are mapped on each chromosome. Three additional markers were mapped to unplaced scaffolds (Table S3).

The addition of more contiguous annelid genomes in the future will reveal the extent of chromosomal rearrangement that has occurred in the annelids, but initial investigation revealed little evidence of macro-synteny across *S. benedicti* and the three other annelid genomes. The genome’s Hox gene cluster was found on chromosome 7 by tBLASTn with queries from other annelids including *C. teleta, Platynereis dumerilii*, and *Myzostoma cirriferum*. This genome contains the full 11 *Antp* set of Hox genes on this chromosome.

## Discussion

The *S. benedicti* genome assembly and annotation provide a critical tool for understanding the genetic basis of phenotypic diversity, including genomic modifications that ultimately lead to evolutionary changes in ontogeny. The *S. benedicti* genome adds to the growing collection of assembled and annotated Lophotrochozoan genomes which collectively can address essential questions of genome and gene function evolution. There are limited full annelid genome assemblies, but *S. benedicti* is one of the most complete and contiguous genomes for the Annelida to date (Table 4). This assembly provides a methodology for other lophotrochozoan genomes that are confounded by limited tissue availability, heterozygosity, and low-representation for lineage-specific genes. The heterozygosity estimated from the Illumina reads is 0.29%, after nine generations of sib-mating, lower than most Lophotrochozoans (Kocot et al. 2020; Varney et al. 2020). Heterozygosity in outbred *S. benedicti* has been estimated at 0.5 to 1% (Rockman 2012) and the modest reduction after inbreeding may reflect the effects of inbreeding depression.

There are some notable assembly caveats: We used females for the Illumina reads because males have a long Y chromosome that is likely repetitive (Zakas et al. 2018), and we wanted to minimize assembly issues. The contents of the Y chromosome, and the sex chromosomes in general, warrants further investigation, especially as a QTL map to the X chromosome and has contrasting directional parental contributions to offspring size (Zakas and Rockman 2020). This genome and transcriptome are generated from planktotrophic animals only, leaving the possibility that structural rearrangements or duplications may have happened between the two types. Previous work has indicated that major genomic rearrangements are unlikely to be a major source of genome divergence (Zakas et al. 2018). Future genomic sequencing and mapping of the lecithotrophic morph should reveal regions of genomic divergence between the types.

This high-quality, chromosomal-scale genome will aid in future population and functional genomics analysis in *S. benedicti*. Genomic comparisons with other Lophotrochozoan animals, as they become increasingly available, will also provide insight into lineage-specific novelty with respect to *poecilogony* and the evolution of marine larval forms. We find that despite having a larger genome than other annelids, and a unique ability to produce different larval types, the *S. benedicti* genome does not contain significant genomic duplications. There are lineage-specific repetitive regions and longer introns and intergenic regions in *S. benedicti* compared to other current annelid genomes.

## Methods

### Animal Collection

We collected animals from Newark Bay, Bayonne, New Jersey, in 2011. To reduce heterozygosity in the animals to be sequenced, we generated an inbred line by mating siblings and crossing a single male and single female each generation. In 2018, we selected adults from the F_9_ generation, flash froze them in liquid nitrogen, and shipped them to Dovetail Genomics (Santa Cruz, CA, USA,) for Illumina shotgun sequencing, Hi-C, and Chicago proximity-ligation sequencing. Prior to freezing, we isolated the animals from food for 24 hours to allow them to purge their guts, and in some cases we dissected their guts away before freezing. For Oxford Nanopore sequencing, we used an additional 71 wild-caught females, collected from the same locality in Bayonne, New Jersey. The genetic map used to scaffold the assembly derives from a cross of a Bayonne female and a male from Long Beach, California, as previously described (Zakas et al. 2018).

### Illumina Shotgun Sequencing

Dovetail Genomics isolated genomic DNA from flash-frozen animals and prepared PCR-free short- and long-insert sequencing libraries. Libraries were sequenced in 2×150bp, yielding 837M read pairs. K-mer analysis of these data suggested a substantial amount of residual heterozygosity (K-mer=31, %SNP heterozygosity= 0.29). As preliminary assembly attempts did not produce sufficient contiguity for scaffolding, we next generated long-read data via Nanopore sequencing (Oxford Nanopore).

### Nanopore Sequencing

For each library, genomic DNA from 20-21 wild caught Bayonne worms was extracted using a CTAB/Phenol Chloroform Isoamyl extraction, yielding ~5ug total gDNA as measured by Qubit DNA kit. Fresh DNA extractions were necessary to ensure HMW DNA as even short-term storage of gDNA at −80C resulted in rapid degradation. We followed the Oxford protocol 1D gDNA selecting for long reads (SQK-LSK109) including the BluePippin size selection step with the 18-27 Kb cassette set to a broad collection range of 11-50 Kb. We used wide-bore low-adhesion tips and minimal pipetting to avoid DNA breakage. Our flowcell and kit combination was (FLO-MIN106D R9 Kit: LSK109). Additional modifications to the Oxford protocol are as follows: Three elutions from the BluePippin cassette returned the most HMW DNA. Elution from the Ampure beads was increased to 15 min at 37°C. The end prep cycle was modified to 20°C for 10 min and 65°C for 10 min. Crucially the adaptor ligation step was extended to 2.5-3 hours at RT. We used 4 flow cells, with 5 library preps (one flowcell was reused).

### Physical Assembly by HiRise

Dovetail generated an initial physical assembly using the Nanopore data with two rounds of polishing by Racon (Vaser et al. 2017). The Nanopore consensus was polished with the Illumina shotgun data. To scaffold this draft into larger components, Dovetail generated Chicago proximity-ligation data (148M 2×150bp read pairs) and Hi-C data (225M 2×150bp read pairs). Using the HiRise software pipeline (Putnam et al. 2016), these data produced an assembly with scaffold N50 of 53.55 Mb.

### Genome Correction with Genetic Map (Chromonomer)

Chromonomer is a tool to combine a scaffolded, unfinished genome sequence with a genetic map by aligning the markers in the QTL map to the genome and re-arranging scaffolds in the unfinished assembly at contig breaks or low-confidence alignment regions such that they match the order of the markers in the map according to their recombination distance. Chromonomer evaluates alignments of each marker in the genetic map to the reference genome and assigns each marker as a node. Where two or more nodes occur in a contiguous sequence, it rearranges the underlying scaffolds to match the genetic map. Out of 702 markers in the genetic map, 598 total markers mapped to the final genome assembly with alignment scores over 10 (Figure 4). In our analysis Chromonomer makes 394 rearrangements, all of which occur at contig breaks where separate contigs were scaffolded together denoted by ‘N’ ambiguity symbols.

### Sequence Decontamination

The genome assembly was examined for potential contaminants via a BlastN (blast+ ver. 2.9.0) search of each of the scaffolds against the NCBI Nucleotide Database reporting back the top ten matches. If all of the top matches were bacterial or viral, the scaffold was flagged as potential contamination. The percent coverage and percent identity of match for each potential contaminant was then assessed to ensure at least an 85% identity between nucleotide sequences and the majority of the scaffold covered. A total of 85 scaffolds were removed from the assembly. The main contaminating species best matched *Erythrobacter flavus*.

### Assembly Quality Assessment

The completeness of the genome assembly was assessed by BUSCO v.3.0.2 (Simão et al. 2015) using the Metazoa odb9 dataset with Augustus species Fly.

### Gene Annotations

For the annotations we generated a PacBio Iso-Seq RNA transcriptome: RNA was extracted (Qiagen) and pooled from a mix of males and females from all stages including embryos. Total RNA was frozen at −80C and sequenced by the Duke Sequencing core, with no size selection on 2 SMRT cells. 264,600 reads were generated. Trimmed reads were clustered and polished with PacBio SMRT Link version 8.0 Isoseq3 tools. There are 24,317 Iso-Seq transcripts. High quality Iso-Seq reads were mapped to the chromosome sequences with GMAP (Wu and Watanabe 2005) version 2020-06-30. Alignment format was set to “gene” and failed alignments were suppressed from the gff3-formatted output.

Gene predictions were generated using maker (Cantarel et al. 2008) version 2.31.10. The 11 chromosomes as well as the 6,101 unplaced scaffolds were used for the predictions. A fasta file of high-quality Iso-Seq sequences (n=24,317) were provided as same-species EST evidence. Protein homology evidence came from 31,978 *Capitella teleta* proteins downloaded from NCBI. Softmasking was selected for repeat masking.

The full set of 41,088 predicted proteins from transcripts were used in a BlastP search of NCBI’s RefSeq protein database with a significance cut-off of 1.0e-05. The GI number for the top match for each of the proteins was extracted and *blastdbcmd* used to assign a descriptor associated with that GI number. A series of scripts were written to create a lookup table which associated a protein name with the protein ID and gene name for each top match. These were added to the annotation file generated by maker. 37,822 (92%) proteins have a putative annotation.

### Repetitive Element Classification

RepeatModeler v2.0.1 (Flynn et al. 2020) was run to identify transposable elements and classify them into families. RepeatMasker was then run using a combination of the Dfam and RepBase databases as well as the species-specific RepeatModeler classifications.

## Availability of data and materials

Results from the genome assembly and annotation are available: http://zakas.statgen.ncsu.edu/jbrowse/?data=sbenedicti

The genome assembly is available at NCBI [BioProject: SUB9470235].

## Acknowledgements

We would like to thank Mohammed Khalfan for assistance with scripts to basecall on the NYU HPC cluster, Julian Catchen for feedback on Chromonomer, and the NCSU bioinformatics consulting core for help with hosting Jbrowser. Thanks to Luke Noble for assistance with early short-read assemblies. Thanks to John Yuen and Arielle Martel for help maintaining laboratory cultures. This work was supported by the Zegar Family Foundation, National Science Foundation (CAREER grant number IOS-1350926 to M.V.R.), National Institute of Health (grant number GM108396 to C.Z.), and startup funds from NYU and NCSU.

