## Supplemental Tables and Figures for "The genome of the poecilogonous annelid *Streblospio benedicti*"

<u>**Supplemental Materials**</u>

**Table of contents**

Figure S1. Link Map From Dovetail
Figure S2. Additional data from OrthoVen2 showing gene counts in each set of clusters.
Table S1. Additional data from OrthoVen2 for four annelid genomes.
Table S2. Enriched Go Categories for gene clusters unique to *C. teleta*(A) and *S. benedicti*(B)
Table S3. Locations for markers mapped to the genome.
Table S4. RepeatMasker output characterizing genomic repeats

**Figure S1. Link Map From Dovetail**

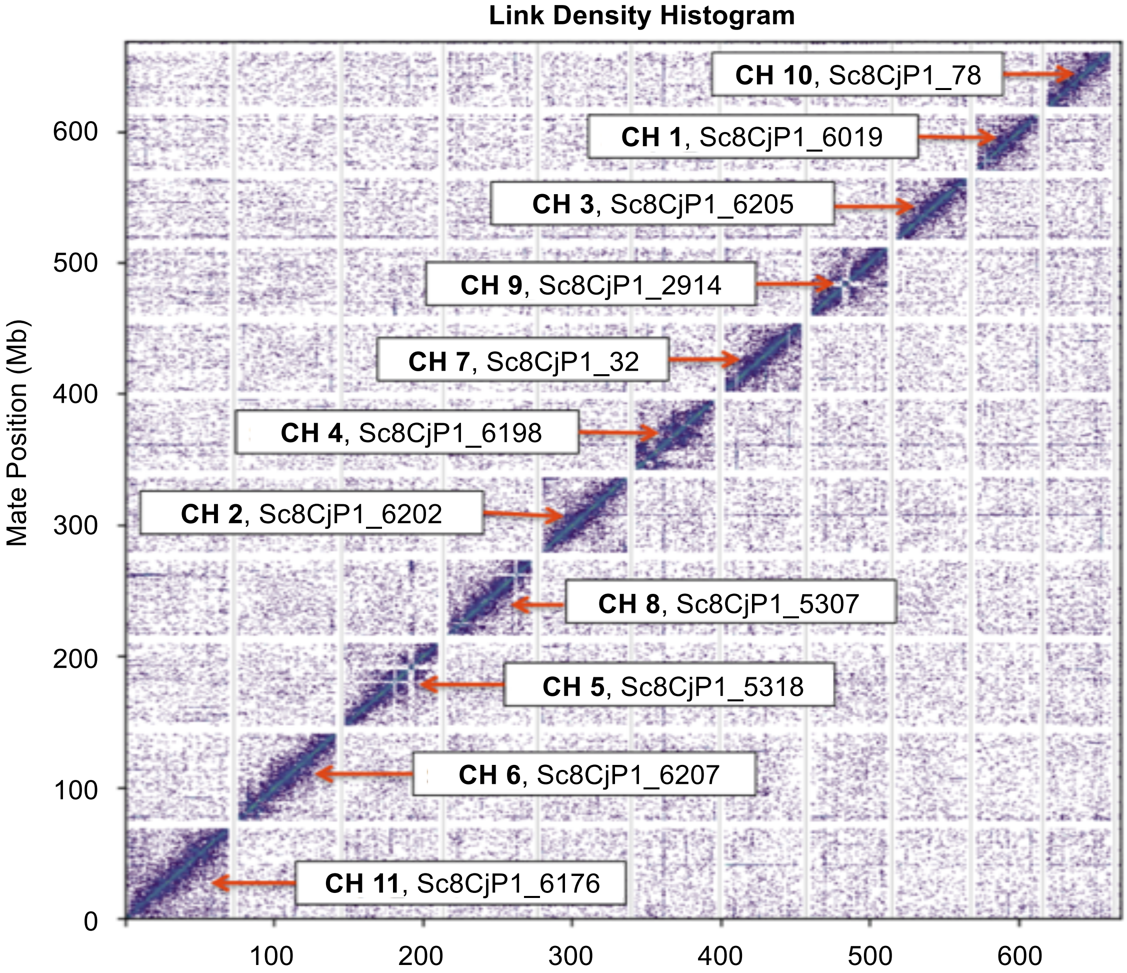

**Figure S2. Additional data from OrthoVen2 showing gene counts in each set of clusters.**

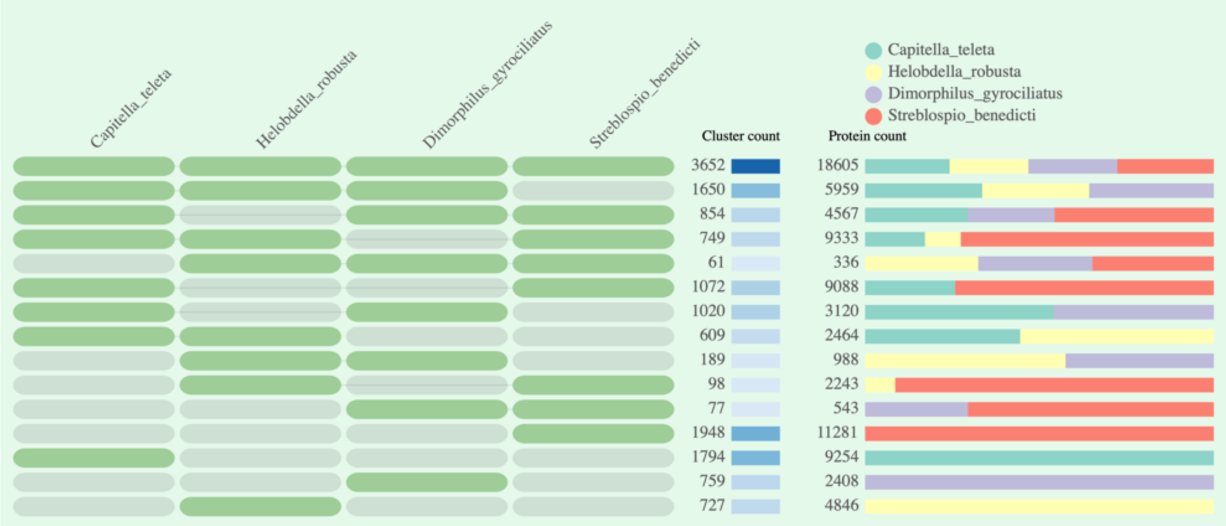


**Table S1. Additional data from OrthoVen2 for four annelid genomes.**
The species form 15,259 clusters, 13,230 orthologous clusters (at least contains two species) and 2029 single-copy (one gene per species) gene clusters.

| Species | Proteins | Clusters | Singletons |
|---|---|---|---|
| *Capitella_teleta* | 32175 | 11400 | 8328 |
| *Helobdella_robusta* | 23432 | 7735 | 9360 |
| *Dimorphilus_gyrociliatus* | 16175 | 8262 | 3639 |
| *Streblospio_benedicti* | 41088 | 8511 | 6508 |

**Table S2. Enriched Go Categories for gene clusters unique to *C. teleta*(A) and *S. benedicti*(B)**

A. *C. teleta* GO enriched categories:

| GO ID | Namespace | Name | Count | p-value |
|---|---|---|---|---|
| GO:0006790 | biological_process | sulfur compound metabolic process | 10 | 1.72E-09 |
| GO:0008484 | molecular_function | sulfuric ester hydrolase activity | 8 | 4.43E-07 |
| GO:0006313 | biological_process | transposition, DNA-mediated | 9 | 8.61E-07 |
| GO:0006486 | biological_process | protein glycosylation | 16 | 1.80E-05 |
| GO:0007204 | biological_process | positive regulation of cytosolic calcium ion concentration | 9 | 4.39E-05 |
| GO:0007585 | biological_process | respiratory gaseous exchange | 4 | 0.000104122 |
| GO:0007165 | biological_process | signal transduction | 30 | 0.000250139 |

B. *S. benedicti* GO enriched categories:

| GO ID | Namespace | Name | Count | p-value |
|---|---|---|---|---|
| GO:0046872 | molecular_function | metal ion binding | 48 | 6.82E-21 |
| GO:0006355 | biological_process | regulation of transcription, DNA-templated | 49 | 1.19E-16 |
| GO:0035282 | biological_process | segmentation | 14 | 1.02E-12 |
| GO:0006486 | biological_process | protein glycosylation | 19 | 5.63E-07 |
| GO:0007165 | biological_process | signal transduction | 38 | 1.00E-06 |
| GO:0045773 | biological_process | positive regulation of axon extension | 6 | 1.03E-06 |
| GO:0010468 | biological_process | regulation of gene expression | 7 | 6.31E-06 |
| GO:0048103 | biological_process | somatic stem cell division | 6 | 1.06E-05 |
| GO:0007596 | biological_process | blood coagulation | 6 | 9.30E-05 |
| GO:0015020 | molecular_function | glucuronosyltransferase activity | 5 | 9.65E-05 |
| GO:0007017 | biological_process | microtubule-based process | 7 | 0.0001059 |
| GO:0060012 | biological_process | synaptic transmission, glycinergic | 8 | 0.0001338 |
| GO:0005524 | molecular_function | ATP binding | 8 | 0.000186 |
| GO:0009409 | biological_process | response to cold | 5 | 0.0002012 |
| GO:0070904 | biological_process | transepithelial L-ascorbic acid transport | 5 | 0.0002012 |
| GO:1901557 | biological_process | response to fenofibrate | 3 | 0.0004007 |
| GO:0009636 | biological_process | response to toxic substance | 6 | 0.0008511 |
| GO:0035202 | biological_process | tracheal pit formation in open tracheal system | 3 | 0.0014851 |
| GO:0000281 | biological_process | mitotic cytokinesis | 3 | 0.0014851 |
| GO:0007160 | biological_process | cell-matrix adhesion | 4 | 0.0015206 |
| GO:0005509 | molecular_function | calcium ion binding | 11 | 0.0018745 |
| GO:0007009 | biological_process | plasma membrane organization | 5 | 0.002183 |
| GO:0019904 | molecular_function | protein domain specific binding | 5 | 0.002183 |
| GO:0048729 | biological_process | tissue morphogenesis | 4 | 0.0025359 |
| GO:0006334 | biological_process | nucleosome assembly | 5 | 0.003034 |
| GO:0007271 | biological_process | synaptic transmission, cholinergic | 8 | 0.0036299 |
| GO:0048771 | biological_process | tissue remodeling | 4 | 0.0039158 |
| GO:0072347 | biological_process | response to anesthetic | 5 | 0.0040889 |

### Table S3. Locations for markers mapped to the genome.

| Chromosome | Position | Marker ID | Chromosome | Position | Marker ID | Chromosome | Position | Marker ID | Chromosome | Position | Marker ID | Chromosome | Position | Marker ID |
|---|---|---|---|---|---|---|---|---|---|---|---|---|---|---|
| 1 | 25260397 | 67 | 6 | 53979595 | 85 | 5 | 34372969 | 899 | 7 | 7558393 | 1087 | 9 | 58224520 | 1244 |
| 7 | 12139825 | 68 | 6 | 38879592 | 667 | 4 | 15448596 | 136 | 5 | 30750381 | 1088 | 7 | 21851946 | 357 |
| 2 | 19156100 | 69 | 5 | 12841187 | 86 | 10 | 3536031 | 901 | 11 | 61373004 | 1090 | 9 | 37330697 | 35 |
| 7 | 31832132 | 517 | 2 | 2390100 | 673 | 10 | 9950047 | 902 | 1 | 662239 | 218 | 7 | 12183862 | 361 |
| 10 | 39546653 | 521 | 5 | 36953425 | 8 | 10 | 9950104 | 903 | 3 | 28439756 | 219 | 7 | 12183949 | 362 |
| 9 | 16754044 | 522 | 6 | 41389374 | 674 | 4 | 23906300 | 904 | 8 | 62827258 | 220 | 9 | 37223475 | 363 |
| 10 | 29714108 | 523 | 9 | 43423585 | 678 | 5 | 50262200 | 913 | 1 | 12832708 | 1093 | 6 | 16957249 | 37 |
| 6 | 19009064 | 70 | 2 | 5609768 | 685 | 7 | 46034391 | 916 | 3 | 15084735 | 223 | 2 | 62616769 | 365 |
| 4 | 13923900 | 525 | 6 | 15410185 | 689 | 9 | 11251935 | 917 | 2 | 21832181 | 225 | 4 | 19404984 | 367 |
| 11 | 60879096 | 530 | 6 | 44505260 | 690 | 5 | 14257376 | 918 | 2 | 5877673 | 226 | 2 | 54317215 | 369 |
| 1 | 11782686 | 534 | 4 | 17528014 | 692 | 4 | 15979927 | 919 | 8 | 18408501 | 229 | 8 | 28161672 | 371 |
| 9 | 29030109 | 536 | 8 | 59054595 | 693 | 10 | 42233607 | 20 | 8 | 16462343 | 230 | 8 | 28161595 | 372 |
| 5 | 52359564 | 537 | 3 | 3056065 | 88 | 2 | 12120222 | 920 | 8 | 16462330 | 231 | 10 | 37299602 | 373 |
| 1 | 3754012 | 538 | 9 | 9277026 | 698 | 9 | 28284833 | 141 | 9 | 45296082 | 1104 | 4 | 7983417 | 375 |
| 10 | 43675750 | 539 | 6 | 15243230 | 700 | 9 | 34078894 | 923 | 10 | 5631764 | 1106 | 2 | 18424825 | 1251 |
| 9 | 17635443 | 540 | 4 | 25233156 | 701 | 6 | 20087233 | 142 | 2 | 31873621 | 1107 | 7 | 26976533 | 378 |
| 9 | 9212772 | 541 | 10 | 12306971 | 702 | 1 | 11808481 | 924 | 4 | 33889187 | 233 | 9 | 41651593 | 1253 |
| 4 | 41215186 | 544 | 4 | 12460785 | 10 | 1 | 11808535 | 925 | 2 | 46709018 | 234 | 4 | 39075100 | 380 |
| 9 | 56561062 | 545 | 4 | 12460866 | 11 | 4 | 4533428 | 928 | 10 | 43533888 | 1110 | 8 | 29566491 | 41 |
| 9 | 11876717 | 72 | 4 | 12460782 | 9 | 11 | 1957386 | 145 | 9 | 11045953 | 1111 | 8 | 54410283 | 381 |
| 11 | 11755029 | 547 | 2 | 42954770 | 706 | 5 | 29322567 | 147 | 8 | 40975268 | 236 | 8 | 40104620 | 384 |
| 4 | 53910595 | 5 | 10 | 796911 | 711 | 10 | 12410851 | 930 | 8 | 64764734 | 237 | 8 | 56240891 | 42 |
| 6 | 38973135 | 549 | 6 | 20296980 | 722 | 9 | 44798763 | 934 | 5 | 13882568 | 1113 | 6 | 30259770 | 392 |
| 6 | 38973168 | 550 | 9 | 139938 | 723 | 8 | 2572809 | 936 | 9 | 10548190 | 1116 | 8 | 50174936 | 1258 |
| 8 | 25381964 | 551 | 3 | 11740304 | 89 | 6 | 45173768 | 937 | 7 | 34093906 | 241 | 7 | 15466033 | 394 |
| 9 | 3357863 | 553 | 5 | 19300732 | 724 | 9 | 40406587 | 944 | 8 | 6811373 | 1118 | 10 | 235293 | 398 |
| 9 | 3302405 | 73 | 8 | 29043900 | 725 | 1 | 14990750 | 945 | 8 | 6811405 | 1119 | 10 | 235275 | 399 |
| 3 | 23768329 | 74 | 7 | 33239645 | 727 | 3 | 41336908 | 946 | 8 | 6811405 | 1120 | 6 | 15156598 | 1259 |
| 3 | 31432567 | 555 | 4 | 8666397 | 90 | 4 | 1849860 | 947 | 8 | 6811417 | 1121 | 9 | 59744862 | 1260 |
| 9 | 13325366 | 556 | 9 | 47649536 | 728 | 4 | 34040799 | 949 | 6 | 19518817 | 243 | 6 | 50543871 | 401 |
| 5 | 30540655 | 557 | 3 | 22507344 | 91 | 9 | 41691338 | 950 | 10 | 4270612 | 245 | 4 | 17345283 | 404 |
| 7 | 90487 | 75 | 2 | 24918710 | 732 | 2 | 27584243 | 151 | 9 | 37580594 | 1126 | 2 | 9383546 | 406 |
| 2 | 44297293 | 76 | 9 | 55260654 | 736 | 8 | 53638435 | 955 | 4 | 8995528 | 1127 | 5 | 45930373 | 1262 |
| 6 | 32678551 | 1310 | 4 | 32445615 | 738 | 4 | 21242036 | 956 | 2 | 13757912 | 247 | 7 | 12006562 | 407 |
| 4 | 6760857 | 1311 | 1 | 6645888 | 740 | 7 | 1218765 | 152 | 8 | 11083978 | 1128 | 9 | 58695843 | 408 |
| 9 | 55135532 | 1314 | 6 | 32305834 | 93 | 7 | 14551620 | 957 | 7 | 36648609 | 1129 | 3 | 47228640 | 409 |
| 9 | 55132055 | 1315 | 2 | 1654480 | 95 | 10 | 16769476 | 958 | 2 | 17490256 | 1130 | 3 | 4445 | 410 |
| 4 | 24153965 | 1316 | 7 | 5922389 | 98 | 4 | 52667193 | 960 | 7 | 22018826 | 250 | 4 | 37765059 | 411 |
| 3 | 9640723 | 1318 | 5 | 17720610 | 742 | 6 | 57561144 | 961 | 5 | 17630180 | 251 | 4 | 17730196 | 412 |
| 8 | 24874263 | 1319 | 9 | 664262 | 743 | 9 | 26788185 | 962 | 4 | 25258258 | 255 | 5 | 34698334 | 416 |
| 8 | 43374242 | 1322 | 8 | 41598302 | 744 | 7 | 5267145 | 963 | 9 | 51223788 | 26 | 6 | 45253043 | 1264 |
| 2 | 35645001 | 1328 | 9 | 28973108 | 748 | 7 | 5267196 | 964 | 8 | 56384180 | 256 | 2 | 17413739 | 418 |
| 7 | 31505417 | 1329 | 1 | 15862219 | 750 | 9 | 5327886 | 966 | 7 | 2782708 | 257 | 3 | 22266583 | 419 |
| 5 | 51274529 | 1331 | 1 | 15862171 | 751 | 2 | 22375916 | 968 | 6 | 4881686 | 1136 | 9 | 35459403 | 1265 |
| 2 | 44516928 | 1334 | 4 | 30616180 | 13 | 1 | 23876783 | 21 | 4 | 26142397 | 262 | 6 | 24421413 | 420 |
| 2 | 44516975 | 1335 | 8 | 36604309 | 99 | 5 | 31164956 | 975 | 6 | 27680190 | 1139 | 6 | 25203205 | 425 |
| 2 | 32636391 | 1336 | 2 | 44578679 | 752 | 5 | 45920989 | 981 | 8 | 40102907 | 267 | 3 | 13485355 | 427 |
| 6 | 39331388 | 1337 | 4 | 37429983 | 100 | 2 | 24469920 | 158 | 1 | 12787707 | 1144 | 2 | 37425961 | 1272 |
| 6 | 30795258 | 1339 | 4 | 46370930 | 101 | 1 | 17070100 | 983 | 6 | 7651250 | 268 | 8 | 37315948 | 431 |
| 9 | 39630398 | 1340 | 4 | 46370927 | 102 | 8 | 62845456 | 984 | 8 | 23363479 | 269 | 8 | 18999137 | 432 |
| 10 | 1103782 | 1341 | 4 | 37853772 | 756 | 7 | 45017526 | 986 | 9 | 5337397 | 270 | 6 | 4680275 | 1274 |
| 8 | 36589776 | 1342 | 7 | 32037729 | 758 | 3 | 22287758 | 991 | 6 | 59681337 | 1151 | 5 | 30072360 | 433 |
| 7 | 33064087 | 1345 | 9 | 12897570 | 103 | 8 | 6499844 | 160 | 7 | 12091922 | 1152 | 9 | 58336625 | 434 |
| 4 | 50997052 | 1346 | 9 | 29584530 | 760 | 6 | 24809682 | 22 | 9 | 35739678 | 272 | 9 | 1551863 | 436 |
| 4 | 50997035 | 1347 | 2 | 27938280 | 761 | 3 | 22898300 | 165 | 7 | 46045181 | 1153 | 4 | 34684664 | 437 |
| 5 | 28967459 | 1348 | 10 | 38414813 | 764 | 4 | 19280729 | 168 | 6 | 41732488 | 1156 | 2 | 24050149 | 438 |
| 9 | 15679866 | 1349 | 9 | 46443229 | 766 | 6 | 59691360 | 1000 | 1 | 17073557 | 275 | 5 | 34327021 | 440 |
| 10 | 19744045 | 1350 | 8 | 162655 | 768 | 6 | 59691354 | 999 | 8 | 6734970 | 277 | 4 | 32571 | 1275 |
| 6 | 41774439 | 1352 | 5 | 40587851 | 105 | 3 | 17203564 | 1002 | 1 | 5930426 | 1158 | 4 | 32601 | 1276 |
| 6 | 51041220 | 1353 | 10 | 16090788 | 772 | 9 | 27642756 | 1003 | 11 | 2980301 | 1160 | 6 | 31651044 | 1277 |
| 1 | 16186025 | 1358 | 2 | 29786623 | 775 | 6 | 4728205 | 1010 | 7 | 23984442 | 279 | 2 | 35964626 | 48 |
| 6 | 5672890 | 1360 | 6 | 64319997 | 776 | 4 | 10409887 | 1011 | 7 | 23984455 | 280 | 7 | 9814888 | 441 |
| 9 | 16407336 | 1361 | 1 | 38138261 | 778 | 4 | 36380760 | 174 | 2 | 41226792 | 1162 | 7 | 9814884 | 442 |
| 8 | 62857851 | 1364 | 3 | 26669387 | 779 | 6 | 44237321 | 175 | 7 | 14184144 | 28 | 7 | 9814833 | 443 |
| 1 | 21469633 | 1367 | 2 | 32935040 | 781 | 10 | 19213245 | 178 | 4 | 15819654 | 1163 | 6 | 23550407 | 49 |
| 9 | 51181691 | 1373 | 5 | 46142545 | 108 | 10 | 19213174 | 179 | 7 | 2864559 | 281 | 8 | 21656330 | 445 |
| 2 | 23073550 | 1375 | 1 | 12456 | 782 | 8 | 21205986 | 1014 | 2 | 43191524 | 283 | 10 | 14477461 | 447 |
| 4 | 16560000 | 77 | 10 | 14155795 | 783 | 4 | 31789286 | 180 | 4 | 21342872 | 284 | 5 | 19286660 | 448 |
| 8 | 35290265 | 6 | 1 | 12834053 | 788 | 6 | 38710193 | 182 | 8 | 33461471 | 287 | 6 | 13460857 | 1282 |
| 1 | 30155911 | 1378 | 9 | 60468670 | 111 | 2 | 17396213 | 183 | 5 | 51091963 | 1168 | 8 | 44428983 | 450 |
| 1 | 30155915 | 1379 | 2 | 28722555 | 790 | 5 | 48032308 | 184 | 6 | 1195572 | 1169 | 7 | 20476932 | 453 |
| 8 | 20491702 | 565 | 5 | 13950152 | 791 | 9 | 5704711 | 1022 | 6 | 1195495 | 1170 | 4 | 39045824 | 454 |
| 9 | 47734390 | 1380 | 4 | 12874879 | 795 | 2 | 24171938 | 1025 | 4 | 52649256 | 292 | 7 | 31394301 | 459 |
| 9 | 47734478 | 1381 | 2 | 25589762 | 796 | 8 | 50171088 | 1030 | 8 | 25485596 | 293 | 3 | 15365491 | 1289 |
| 11 | 9657715 | 7 | 6 | 54674820 | 113 | 5 | 12348065 | 1034 | 5 | 43012133 | 1175 | 2 | 7284998 | 462 |
| 8 | 13313647 | 1382 | 8 | 33607283 | 798 | 1 | 34104063 | 185 | 8 | 56410300 | 1181 | 9 | 43513167 | 463 |
| 1 | 21854299 | 566 | 8 | 33607359 | 799 | 3 | 35025157 | 1035 | 3 | 37551517 | 1183 | 2 | 2094530 | 1293 |
| 7 | 4052380 | 1385 | 4 | 14025528 | 800 | 6 | 56267972 | 1036 | 3 | 46528152 | 1184 | 2 | 2094587 | 1294 |
| 7 | 5355807 | 569 | 1 | 26696172 | 806 | 8 | 42859488 | 1037 | 5 | 43044667 | 297 | 10 | 19257458 | 51 |
| 8 | 32690903 | 571 | 1 | 26696126 | 807 | 8 | 42859458 | 1038 | 9 | 43356260 | 1190 | 8 | 42476922 | 1296 |
| 5 | 36983496 | 1386 | 9 | 38138157 | 808 | 4 | 41223168 | 1039 | 11 | 22559723 | 1193 | 1 | 21884917 | 467 |
| 1 | 37941552 | 1389 | 7 | 14106440 | 810 | 3 | 38731678 | 1040 | 6 | 9713053 | 299 | 4 | 37256729 | 468 |
| 5 | 5821346 | 574 | 11 | 16958808 | 813 | 3 | 20354184 | 1041 | 10 | 15769974 | 1194 | 4 | 16269341 | 469 |
| 8 | 26732318 | 576 | 8 | 17150792 | 814 | 10 | 32850027 | 1043 | 3 | 29512179 | 301 | 5 | 30362506 | 472 |
| 4 | 31015566 | 578 | 8 | 6451277 | 114 | 2 | 21777541 | 1044 | 2 | 10256363 | 302 | 9 | 14832333 | 473 |
| 4 | 45459957 | 580 | 7 | 36238265 | 824 | 4 | 39983442 | 1046 | 5 | 34678221 | 1195 | 2 | 46467370 | 53 |
| 11 | 16554180 | 581 | 4 | 35843837 | 115 | 9 | 60478841 | 189 | 1 | 12783879 | 305 | 7 | 19429085 | 475 |
| 5 | 23695227 | 584 | 9 | 32521709 | 116 | 9 | 60479360 | 190 | 2 | 17541934 | 306 | 8 | 38302608 | 476 |
| 3 | 18382131 | 585 | 6 | 57818365 | 119 | 9 | 60479378 | 191 | 9 | 41602989 | 1203 | 2 | 17519961 | 478 |
| 2 | 22918953 | 593 | 2 | 46196198 | 829 | 9 | 60485539 | 192 | 7 | 34230086 | 312 | 2 | 17516901 | 479 |
| 7 | 25041681 | 594 | 3 | 17604960 | 832 | 9 | 7283447 | 1050 | 7 | 34230122 | 313 | 2 | 17513995 | 482 |
| 2 | 27243959 | 597 | 4 | 37269418 | 834 | 9 | 13463943 | 1051 | 2 | 22473997 | 1204 | 2 | 17512838 | 483 |
| 10 | 43551791 | 79 | 7 | 39744582 | 839 | 2 | 27856962 | 1055 | 1 | 10951907 | 1207 | 2 | 17511431 | 485 |
| 10 | 43551741 | 81 | 10 | 30011429 | 845 | 2 | 40208835 | 194 | 7 | 23643707 | 314 | 2 | 17508788 | 487 |
| 5 | 7381427 | 599 | 4 | 45175452 | 847 | 3 | 23821776 | 196 | 7 | 42574759 | 315 | 2 | 17507809 | 488 |
| 2 | 43707484 | 605 | 11 | 58977320 | 848 | 5 | 711230 | 1061 | 7 | 25456741 | 1212 | 2 | 17506696 | 490 |
| 2 | 43707496 | 606 | 9 | 2309021 | 850 | 5 | 711224 | 1062 | 9 | 48915550 | 317 | 2 | 17505609 | 491 |
| 5 | 51293375 | 607 | 7 | 4083664 | 122 | 2 | 46934871 | 198 | 4 | 17426537 | 318 | 2 | 17498691 | 493 |
| 5 | 51293421 | 608 | 6 | 64858980 | 124 | 2 | 46934978 | 199 | 1 | 93611 | 320 | 1 | 9303085 | 57 |
| 8 | 31351397 | 609 | 10 | 41037109 | 859 | 9 | 9526479 | 25 | 9 | 44564427 | 322 | 11 | 56782520 | 494 |
| 8 | 31351332 | 610 | 8 | 29214989 | 860 | 6 | 25048043 | 200 | 9 | 41847672 | 324 | 6 | 39049455 | 495 |
| 3 | 13198955 | 611 | 1 | 20342093 | 863 | 6 | 28916334 | 1063 | 8 | 17156771 | 31 | 7 | 20639047 | 496 |
| 6 | 28297090 | 612 | 11 | 10137430 | 865 | 10 | 37815896 | 1065 | 7 | 21887888 | 1215 | 3 | 29025887 | 497 |
| 8 | 7613491 | 613 | 8 | 63146302 | 870 | 7 | 36867339 | 202 | 9 | 47014185 | 328 | 11 | 57721622 | 1298 |
| 11 | 25034460 | 616 | 3 | 37292578 | 871 | 6 | 51346307 | 1068 | 9 | 10567988 | 331 | 8 | 36045333 | 498 |
| 11 | 16916645 | 618 | 9 | 58748192 | 126 | 4 | 21171130 | 1069 | 9 | 15370007 | 1221 | 6 | 43114092 | 501 |
| 4 | 41074882 | 620 | 4 | 26423160 | 875 | 4 | 40122925 | 1070 | 4 | 50926587 | 334 | 4 | 31639234 | 504 |
| 4 | 26161452 | 622 | 7 | 40240149 | 127 | 7 | 7918831 | 204 | 4 | 50926596 | 335 | 4 | 23720090 | 505 |
| 4 | 26161438 | 623 | 3 | 37515329 | 876 | 9 | 52980516 | 205 | 4 | 17123859 | 336 | 4 | 23775521 | 507 |
| 9 | 47578933 | 83 | 6 | 29673495 | 128 | 5 | 42120466 | 1072 | 1 | 17009483 | 32 | 6 | 45970146 | 509 |
| 8 | 29047803 | 633 | 6 | 27790081 | 882 | 11 | 47661861 | 1073 | 10 | 32982614 | 1223 | 5 | 51009333 | 511 |
| 9 | 33709949 | 639 | 4 | 23111643 | 885 | 4 | 32826582 | 207 | 4 | 41714296 | 1225 | 3 | 28332996 | 513 |
| 10 | 9638917 | 641 | 8 | 40265287 | 887 | 4 | 28852086 | 1079 | 8 | 28075195 | 344 | 9 | 11169771 | 64 |
| 10 | 9638929 | 642 | 1 | 10421710 | 890 | 2 | 2043065 | 1080 | 6 | 52629042 | 1234 | 7 | 24256594 | 515 |
| 8 | 14231438 | 643 | 11 | 13887753 | 891 | 5 | 2974075 | 1081 | 7 | 8421446 | 1237 | 8 | 56732729 | 516 |
| 3 | 23682964 | 652 | 9 | 3405113 | 130 | 11 | 19386639 | 1083 | 8 | 26505547 | 1239 | Scaffold_541 | 541229 | 703 |
| 6 | 36016717 | 655 | 9 | 22476238 | 893 | 8 | 37001647 | 214 | 8 | 26505505 | 1240 | Scaffold_184 | 19314 | 177 |
| 10 | 38834002 | 657 | 9 | 6392487 | 132 | 8 | 46707187 | 1084 | 1 | 30233888 | 1241 | Scaffold_397 | 9004 | 1172 |
| 8 | 12143637 | 660 | 2 | 49881664 | 894 | 2 | 8390317 | 216 | 6 | 45140548 | 350 | | | |
| 8 | 12140806 | 661 | 2 | 11377404 | 896 | 6 | 46685714 | 217 | 7 | 9029298 | 351 | | | |

**Table S4. RepeatMasker output characterizing genomic repeats**

==================================================

| | |
|---|---|
| sequences: | 6112 |
| total length: | 701454808 bp |
| GC level: | 37.93 % |
| bases masked: | 283086843 bp ( 40.36 %) |

==================================================

| | number of elements* | length occupied | percentage of sequence |
|---|---|---|---|
| SINEs: | 28628 | 5231960 bp | 0.75 % |
| ALUs | 0 | 0 bp | 0.00 % |
| MIRs | 0 | 0 bp | 0.00 % |
| LINEs: | 92268 | 29141093 bp | 4.15 % |
| LINE1 | 716 | 96824 bp | 0.01 % |
| LINE2 | 13068 | 3635866 bp | 0.52 % |
| L3/CR1 | 12084 | 4436605 bp | 0.63 % |
| LTR elements: | 27577 | 15559465 bp | 2.22 % |
| ERVL | 0 | 0 bp | 0.00 % |
| ERVL-MaLRs | 0 | 0 bp | 0.00 % |
| ERV_classI | 663 | 98352 bp | 0.01 % |
| ERV_classII | 166 | 12815 bp | 0.00 % |
| DNA elements: | 38708 | 12966888 bp | 1.85 % |
| hAT-Charlie | 0 | 0 bp | 0.00 % |
| TcMar-Tigger | 0 | 0 bp | 0.00 % |
| **Unclassified:** | **1038899** | **211774011 bp** | **30.19 %** |
| Total interspersed repeats: | | 274673417 bp | 39.16 % |
| Small RNA: | 298 | 47093 bp | 0.01 % |
| Satellites: | 564 | 110176 bp | 0.02 % |
| Simple repeats: | 131118 | 743601 bp | 1.10 % |
| Low complexity: | 17440 | 78030 bp | 0.13 % |

==================================================
